# Factors contributing to the disease ecology of brown crab (*Cancer pagurus*) in a temperate marine protected area

**DOI:** 10.1101/2020.12.14.416628

**Authors:** Charlotte E. Davies, Andrew F. Johnson, Emma C. Wootton, Spencer J. Greenwood, K. Fraser Clark, Claire L. Vogan, Andrew F. Rowley

## Abstract

Marine ecosystems are affected by multiple, well-known stressors like fishing and climate change, but a less documented concern is disease. Marine reserves have been successful in replenishing stocks and aiding recruitment but studies have shown that high population abundances in marine reserves may lead to unwanted secondary effects such as increase in predators and competition, altering trophic webs, and disease. Here, we investigate factors contributing to disease prevalence in a brown crab (*Cancer pagurus*) population around Lundy Island (the UK’s first MPA) after 7 years of no-take protection. Population parameters (size, sex, and abundance), disease (shell disease, *Hematodinium* spp. infection) and injury presence (a known precursor to some disease conditions) were assessed over two years in both fished and unfished areas of the MPA. We found no significant difference in prevalence between the disease prevalence in fished and unfished areas, however overall, the number of injured crabs increased significantly over the two years (12%), as did the prevalence of shell disease (15%). The probability of crabs having shell disease increased significantly in male crabs, and in those missing limbs. The probability of crabs being injured increased significantly in crabs below the minimum landing size. In terms of population parameters, crabs were more prevalent in the fished area compared to the unfished area, thought to be a result of an increase in the predatory European lobster. The findings of the present study highlight potential secondary community changes as a result of MPA implementation. Therefore, surveillance for such changes, as part of MPA management, would provide useful information on the health and overall function of the protected ecosystem.

## INTRODUCTION

Numerous commercial fish stocks have been overexploited and many continue to be fished at unsustainable levels (Jackson et al. 2001, Steadman et al. 2014). This is leading to both a loss of biodiversity and significant concerns for global food security (Pauly et al. 2005, Worm et al. 2009). Conservation areas in marine ecosystems, such as marine protected areas (MPAs) are one management tool that can be used to help reduce fished species decline, aid in stock replenishment and conserve habitats of special interest (Halpern & Warner, 2002, Aburto-Oropeza et al. 2011).

Closure of areas to fishing may, however, result in secondary effects such as overpopulation of a species, in turn altering natural local community compositions (Leo & Micheli 2015, Wood & Lafferty 2015). In some protected areas, changes in species assemblages and population abundance have been shown to have negative effects due to overcrowding such as disease increase, reduction in habitats and change in food web structure due to changes in competition or predation rates (Lebarbenchon et al. 2006, Wootton et al. 2012, Wood et al. 2013, Christianen et al. 2014). With increased abundance comes an increase in both conspecific (within species) and heterospecific (between species) interactions, which can result in injury and limb loss in individuals, contributing to disease in some species (Davies et al. 2015). Disease can influence community composition, age distributions, trophic interactions and biotic structure within a population (Harvell et al. 2002). In some cases, disease is also thought to be exacerbated by the emerging threat of climate change (Burge et al. 2014, Maynard et al. 2016) and can be a useful indicator of ecosystem health (Harvell et al. 1999). The prevalence and distribution of pathogens and disease in marine ecosystems is growing globally (Ward & Lafferty 2004), has been reported across a wide range of taxa over the past three decades (Harvell et al. 2004) and is seen as an often neglected, but emerging field (Lafferty & Hofmann 2016).

*Cancer pagurus*, the edible or brown crab, is an important European fisheries species, with global landings of over 32,000 tonnes in 2014. This species is, however, susceptible to a range of pathogens (Stentiford 2008), the most documented of which, that have been known to cause significant economic and population losses, are pink crab disease and ‘black spot’ or shell disease. Pink crab disease, in some species referred to as bitter crab disease, is caused by the endoparasitic dinoflagellate, *Hematodinium* spp., named as such because of the hyperpigmentation and bitter taste exhibited by some heavily infected species (Wilhelm & Mialhe 1996, Stentiford et al. 2002, Ryazanova 2008). Chatton & Poisson (1931) first reported the disease in France in both harbour *Liocarcinus depurator* and shore crabs *Carcinus maenas*. It has since been found to infect over 40 species of decapod crustaceans worldwide, and because infected animals become unmarketable due to poor muscle quality, *Hematodinium* spp. infections have had huge economic impacts on commercial fisheries (Field et al. 1992, Wilhelm & Mialhe 1996, Shields et al. 2005, Stentiford & Shields 2005).

‘Black spot’ or ‘Burn Spot’ disease, herein referred to as shell disease, is characterised by melanised lesions that can progress to erode through the carapace, exposing underlying soft tissues in the infected individual (Vogan et al. 2008). Mortality occurs due to secondary infection by opportunistic bacteria (Baross et al. 1978, Vogan & Rowley 2002), or during a moult if old and new shells adhere at the lesion site (Fisher et al. 1978). Shell disease has become prevalent in many UK *C. pagurus* and European lobster *Homarus gammarus* fisheries since it was first reported by Pearson (1908). The Isle of Man (UK) fishery displayed shell disease in almost 25% of *C. pagurus* sampled during the summer of 2012 (King et al. 2014), and in a South Wales (UK) fishery, over 50% of *C. pagurus* were affected in 1997-1998 (Vogan et al. 1999). Between 1985-1987, the West coast of Scotland fishery displayed shell disease in almost 100% of individuals, significantly higher than in other crab species analysed in the same survey (Comely & Ansell 1989). Ayres and Edwards (1982) reported 5-7% of affected animals were rejected from a South West Irish fishery and suggested that the incidence of shell disease was higher in lightly fished populations than in established fisheries, where intensive exploitation results in the removal of larger, older crabs from the stock.

The aim of the present study was to examine factors contributing towards these two diseases in a population of *C. pagurus* in both a fished and un-fished area of the Lundy Island MPA in the Celtic Sea, UK. In the past, this MPA has been reported to have higher levels of disease in the unfished area due to increased population density, or overcrowding (Wootton et al. 2012, Davies et al. 2015). Therefore, it was first hypothesized that the population abundance of *C. pagurus* would be higher in the un-fished area of the MPA compared to the fished area, as observed in a previous study (Hoskin et al. 2011). The second hypothesis was that the crabs from the un-fished area would have an increased probability of injury, limb loss and therefore disease than individuals from the fished area due to overcrowding. Individuals missing one or more limbs, or with open wounds are expected to have a higher probability of disease than those with limbs intact, as limb loss creates a large wound, which can act as an aperture for pathogen entry. Our third hypothesis was that, as experienced in previous studies, the unfished area would have a higher prevalence of large individuals (herein classified as individuals over the minimum landing size; MLS ≥ 160 mm carapace width for males and ≥ 140mm for females) which would in turn exhibit an increased probability of disease and injury than smaller individuals (those under MLS) as experienced for some infections by Bateman et al. (2011). In larger animals, longer inter-moult periods give disease more time to manifest and competition brought on by sexual maturity has also been shown to increase in older, larger crustacean species (Edwards 1966).

## MATERIALS AND METHODS

### Study area

The study took place around Lundy Island, 12 miles off the coast of North Devon, England, UK (Fig. 1), which was Britain’s first MPA. Lundy first consisted of a Refuge Zone (RZ), established in 1986 when it was designated as Britain’s first and only Marine Nature Reserve. Here, pot fisheries were authorised, but trawl and net fisheries were prohibited. In 2003, A No-Take Zone (NTZ) was incorporated, where all fishing, including potting, and removal of wildlife is forbidden. Lundy Island became Britain’s first Marine Conservation Zone in 2010.

**Figure 1.**
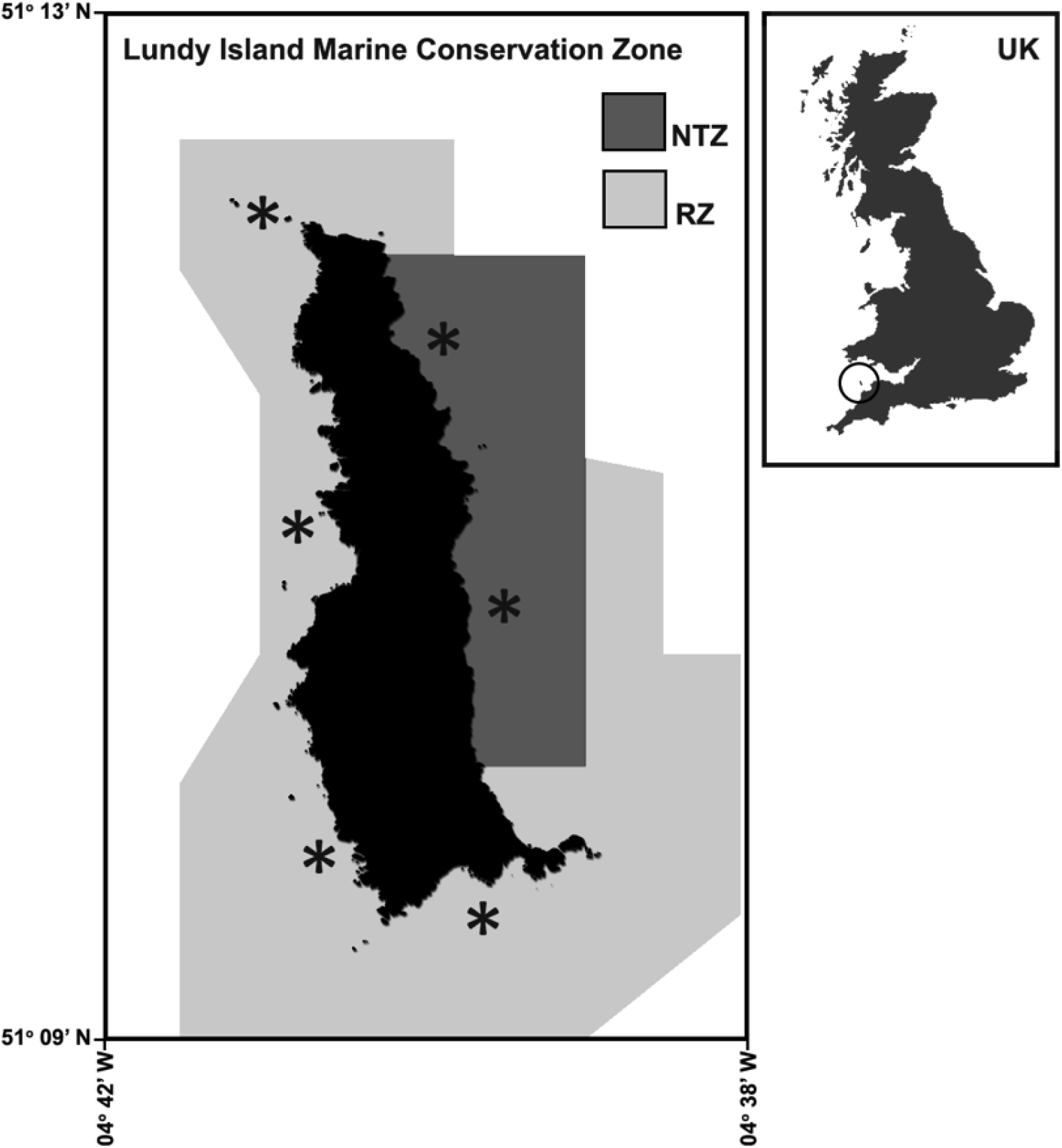
Map showing the Lundy Island MPA Marine Conservation Zone and sampling sites (*) at which strings of pots were deployed within the NTZ and RZ. Lundy’s position relative to the UK is circled in the smaller map. Adapted from Davies *et al.* (2015).

### Population sampling

The *C. pagurus* population around Lundy Island was sampled in May and July 2010, and August 2011. One string of baited commercial parlour pots (35 pots with escape gaps closed) was deployed at a total of 6 sampling sites (4 in the fished RZ and 2 in the unfished NTZ, Fig. 1). In total, 30 strings were deployed in 2010 and 18 in 2011 with a total of 397 crabs sampled (213 in 2010 and 184 in 2011). Each string was similarly baited, immersed for 24h, retrieved and emptied of all catch. Seven measurements were recorded for each *C. pagurus* caught (Table 1). In order to assess the presence of *Hematodinium* spp. using molecular methods, *ca.* 500-700 μl of haemolymph was drawn into 1 ml of 100% analytical grade ethanol using 23 g needles and 2 ml syringes. Crabs exhibiting exoskeletal abnormalities or severe shell disease were photographed (see Figure 2 for examples of these conditions). All individuals were measured, traps were re-baited, re-deployed and all catch returned to the water.

**Table 1.** Parameters recorded for each individual *C. pagurus* caught. Parameters used as predictor variables in binomial logistic regression models are denoted as: *for shell disease, †for *Hematodinium* spp. and ‡for injury.

| Measurement | Description | Measure |
| --- | --- | --- |
| **Berried** | Presence of eggs attached to underside of abdomen of females | Presence vs. Absence (P vs. A) |
| **Carapace width (CW)** | Width across the dorsal carapace at widest point | Continuous measure (mm) |
| **Injury \*,†** | Wounds such as punctures and stress fractures breaching the cuticle. Injuries inflicted during captivity within the pot (i.e. recent, non-melanised breach of the cuticle) were not recorded | P vs. A |
| **Limb loss \*,†** | Missing cheliped or walking leg | P vs. A |
| **Minimum landing size (MLS) \*,†,‡** | MLS (width of carapace) as designated by The Devon and Severn IFCA District shore Fisheries and Conservation Authority byelaw (≥140mm for females, ≥160mm for males) | 0 (below MLS) or 1 (above MLS) |
| **Sex \*,†,‡** | Gender of crab (male or female) | M vs. F |
| **Shell disease presence** | Shell disease (photographic examples in Figure 2) | P vs. A |

**Figure 2.**
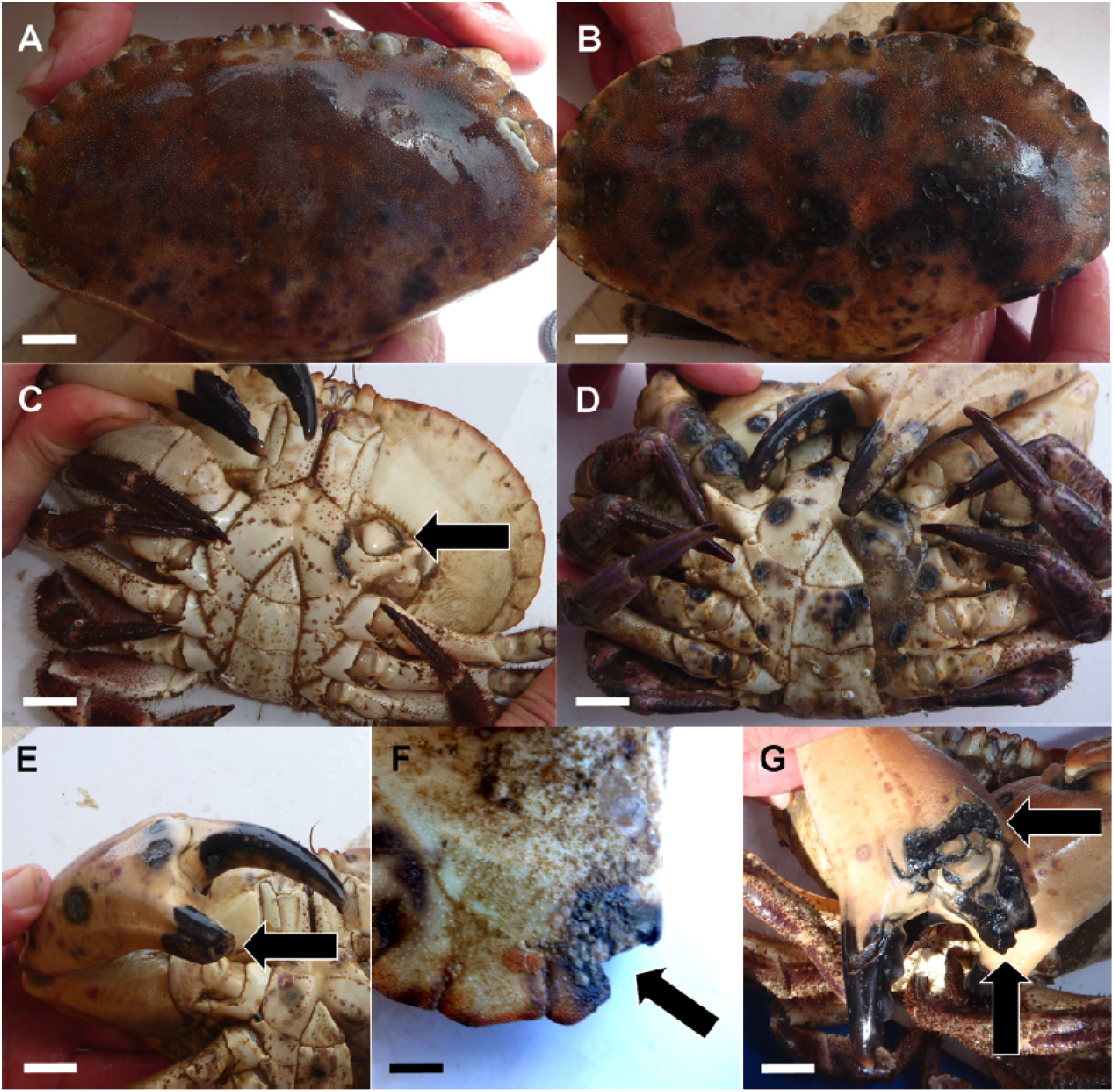
Examples of *C. pagurus* displaying a healthy (A) and shell diseased (B) dorsal carapace, a healthy (C) ventral carapace with limb loss (black arrow), a shell diseased ventral carapace (D) and (E-G) examples of injury at various sites (black arrows): (E) the propodus region of a chela, missing, (F) edge of carapace and, (G) the propodus region of a chela. Scale bars represent 2 cm.

**Figure 3.**
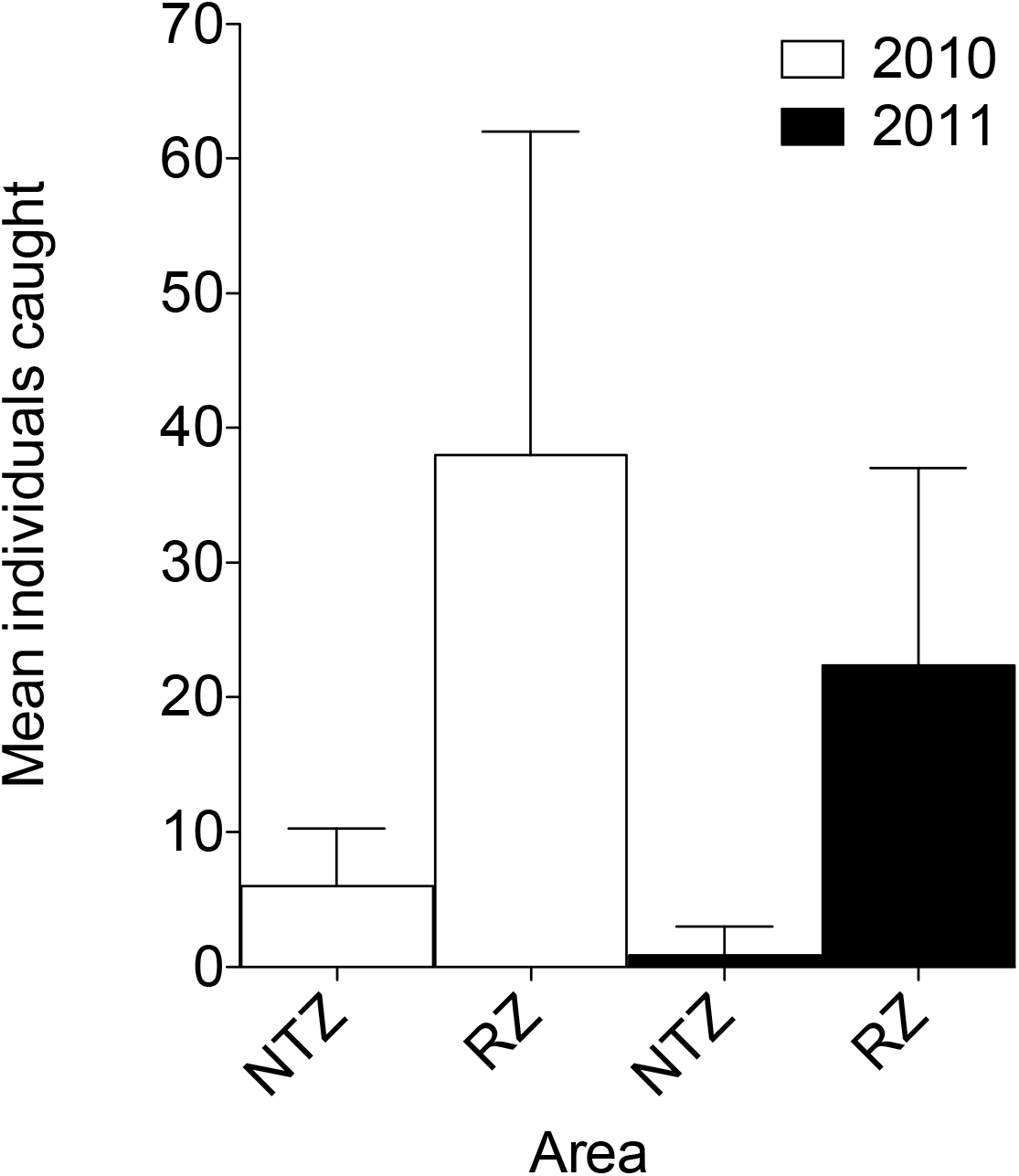
Mean number of edible crabs (with 95% CI) caught per 35-pot string in the No-Take Zone (NTZ) and Refuge Zone (RZ) in 2010 and 2011.

### Surveillance of haemolymph pathogen communities

#### DNA extraction

DNA was extracted from haemolymph using a modified version of Ivanova et al. (2006, see Section S1, Supplementary Materials). DNA was eluted with water, stabilized with Tris-EDTA buffer (10 X) and used as the template for polymerase chain reactions (PCR). Haemolymph DNA extraction was optimized to ensure detection of *Hematodinium* spp. by using known, positive controls initially derived from *C. pagurus*, confirmed by sequencing.

#### PCR conditions

All PCR were carried out using primers synthesized by Eurofins MWG Operon (Ebersberg, Germany) and performed on a Bio-Rad PTC-100 Peltier Thermal Cycler before being visualized on a 1.5% agarose gel. First, decapod-specific primers were used to verify the quality of the extracted DNA and the integrity of the PCR reaction (Section S2, Supplementary Materials). DNA was then amplified using *Hematodinium* spp. specific primers optimized by Hamilton et al. (2009) in order to test haemolymph DNA for the presence of *Hematodinium* spp. infection (Section S3, Supplementary Materials). *Hematodinium* spp. positive samples were repeated, the PCR product cleaned up using the Wizard SV Gel and PCR Clean-Up System (Promega, Madison, USA) and sequenced by Eurofins MWG Operon (Ebersberg, Germany). Contigs from sequences were created using the CAP3 sequence assembly programme (Huang & Madan 1999) and identity confirmed using matched positive controls via NCBI BLAST.

### Statistical Analysis

#### Dataset Determination

Data from May and July 2010 were pooled so that one coherent year was used to compare with the 2011 catch data (see Table S1, Supplementary Materials). To minimize the possibility that individuals from May and July sampling trips were not double sampled (i.e. released in May, recaptured in July) each individual in the database was given a unique identifier based on the predictor variables in Table 1 and any individuals sharing the identifier were noted. The number of potential recaptures was 11.61%. In order that individuals were not double sampled on consecutive sample days within each month (i.e. recaptured after day one of a survey and considered a unique individual), the same method was used and 0.36% of individuals were classified as potential recaptures and removed.

#### Population Ecology

Population distributions of males and females were visualised in GraphPad Prism 5.0, plotted per site and year. In order to compare catch and size-frequency data between the NTZ and RZ, data were first tested to follow a normal distribution (using two-sample Kolmogorov-Smirnov tests) followed by either an independent T test (if data was normal) or Mann-Whitney test (if data did not follow a normal distribution). Tests were two-tailed and used a significance level of 0.05. Catch Per Unit of fishing Effort (CPUE) was calculated as the mean number of animals per pot. A linear regression was used to examine the relationship between the CPUE of *C. pagurus* and European lobster, *H. gammarus*, the two commercially viable species found in the pots.

#### Disease and Injury Ecology

Binomial logistic regression models with Logit link functions (following Bernoulli distributions) were used (MASS library, R Development Core Team 2014) to determine whether specific predictor variables (Table 1) had a significant effect on the presence of shell disease, *Hematodinium* spp. infection and injury presence in the crab population sampled. The information theoretic approach was used for model selection and assessment of performance (Richards 2005). To begin, a suite of models ranging from fully additive to models combining all possible combinations of single, interactive terms were generated using the predictor variables highlighted in Table 1. The best models from each suite were selected based on Akaike’s Information Criterion (AIC) which measures model “quality” based on the goodness of fit and parsimony of the model: the lower the AIC, the better the model (Burnham & Anderson 1998, Zuur et al. 2009). Selected initial models are herein referred to as the *full models*. Once selected, each non-significant predictor variable from the full models was sequentially removed using the drop1 function (in R) to produce final models with increased predictive power, herein referred to as the *reduced models*. The drop1 function compares the initial full model with the same model, minus the least significant predictor variable. If the reduced model is significantly different from the initial full model (in the case of binomial response variables, a Chi-squared test is used to compare the residual sum of squares of both the models), then the removed predictor variable is kept out of the new, reduced model. This process continues hierarchically until a final reduced model is produced (Zuur et al. 2009). Fitted probability plots were used to visualize the significant relationships inferred from the reduced models using carapace width (CW) as the independent variable. The probability of a crab having shell disease, *Hematodinium* spp. infection or being injured, was calculated using the following equation:

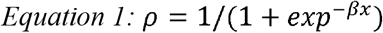

Where ρ is the probability of shell disease, *Hematodinium* spp. infection or injury presence and β*x* is the estimate for the predictor variable analysed (Table 1).

## RESULTS

### General population ecology

Comparative analysis between the two sites (fished; RZ vs. un-fished; NTZ) revealed that there were significantly more crabs caught per string in the RZ than in the NTZ in both the 2010 and 2011 surveys (2010: P = 0.003, t = 3.62, NTZ = 6 ± 1.86, RZ = 38 ± 9.82; 2011: P = 0.002, t = 2.97, NTZ = 0.83 ± 0.83, RZ = 22.38 ± 6.19 [mean values ± SEM], Fig. The CPUE was 6.3 times greater in the RZ than in the NTZ in 2010 and 26.9 times greater in 2011.

There was no significant difference in the number of crabs above the MLS caught between the NTZ and RZ in either 2010 or 2011(P > 0.05, see Table 2 and Fig. 5A-D). Not accounting for the sex of crabs, the size frequency distributions were significantly different between the NTZ and RZ in 2010 (P = 0.005) and 2011 (P < 0.001). There was, however, no significant difference between the size of crabs in the NTZ and the RZ in 2010 (P = 0.205) or 2011 (P = 0.077), even when separating crabs by sex in 2010 (males P = 0.249, females P = 0.488, Fig. 5A-D). Due to the low abundance of crabs caught in the NTZ in 2011 (n male = 4, n female, = 1), size differences between zones could not be tested. No ovigerous (‘berried’) females were caught in either year from either zone.

**Table 2.**
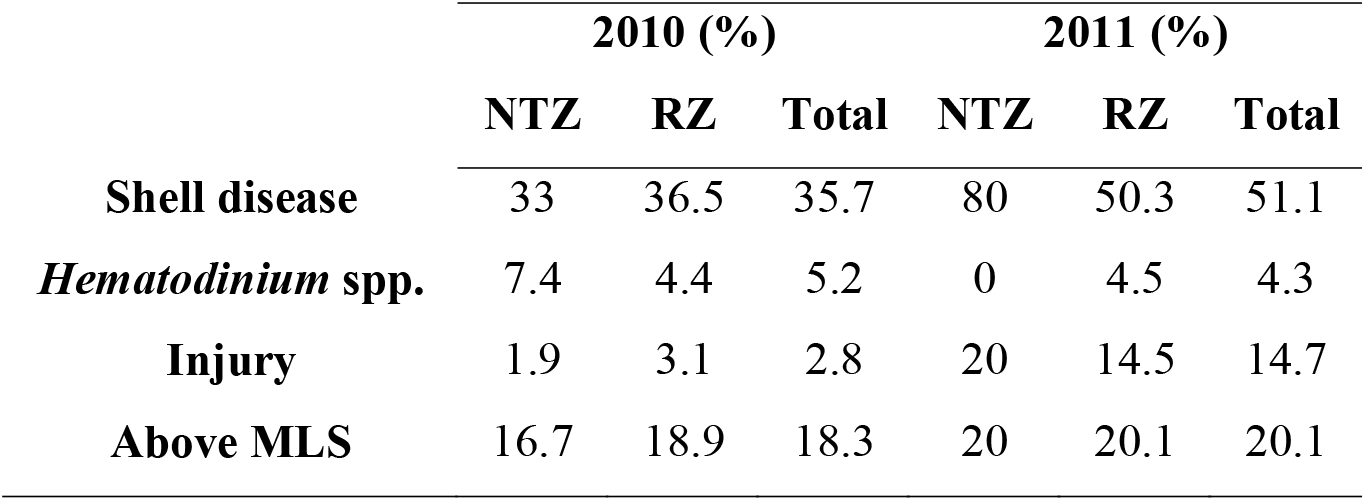
Percentages of crabs according to predictor variable used in models.

### Shell Disease

Comparative analysis between the two sites (fished; RZ vs. un-fished; NTZ) revealed that there was no significant change in shell disease prevalence between sites. However, when combining data from both sites, analysis highlighted that the percentage of shell diseased crabs increased significantly by *ca*.15% between 2010 and 2011 (P = 0.002, Table 2). There was no significant effect of site, limb loss, landing size, injury, sex or *Hematodinium* spp. infection in explaining shell disease presence in 2010 (Table S2, Model S1). Sex, limb loss, and the interaction between *Hematodinium* spp. and sex (infected male crabs), all had a significant effect on the presence of shell disease in 2011 (Table 3). If a crab was male, or a crab was missing a limb, the probability of it having shell disease increased by 87% and 74% respectively (Fig. 6). Crabs which were male and harboured *Hematodinium* spp. were 2% **less** likely to exhibit shell disease.

**Table 3.**
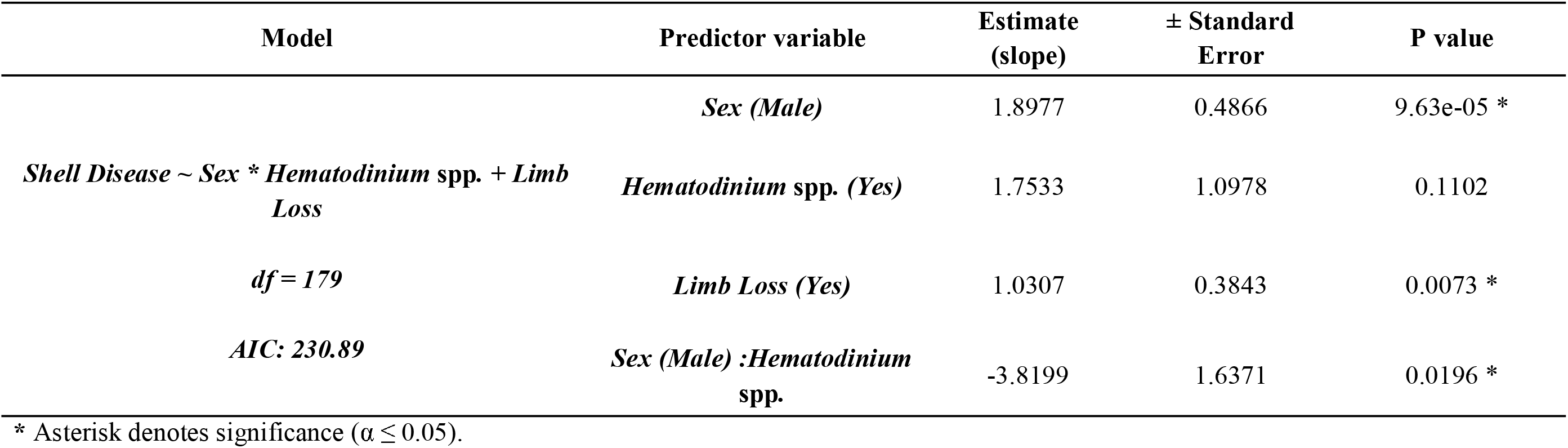
Binomial logistic regression Model 1, reduced from the full model (Table S2, Model S2), testing the effects of sex, *Hematodinium* spp. presence and limb loss on the presence of shell disease in 2011.

### Presence of *Hematodinium* spp

Comparative analysis between the two sites (fished; RZ vs. un-fished; NTZ) revealed that there was no significant change in *Hematodinium* spp. prevalence between sites. However, only 19 out of 397 individuals tested positively for *Hematodinium* spp. (Table 2). None of the predictor variables were significant in predicting the presence of *Hematodinium* spp. in 2010 (Table S2, Model S3). Sex had a significant effect in determining *Hematodinium* spp. infection in 2011 (Table S2, Model S4) but this proved marginally non-significant following model reduction (Table 4).

**Table 4.**
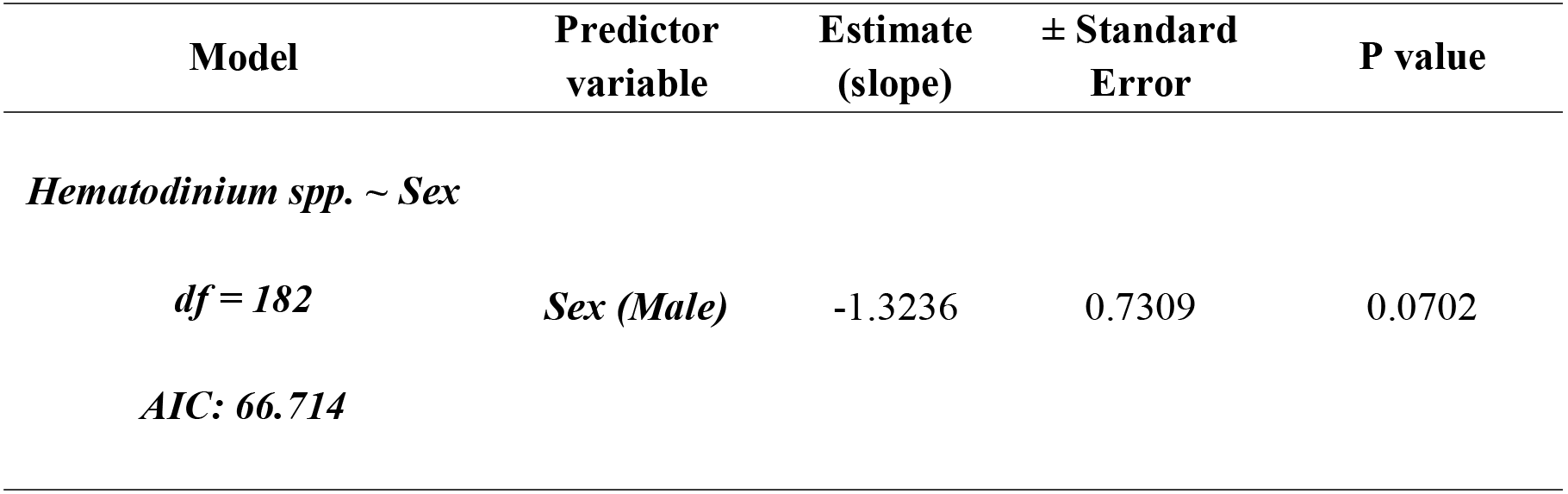
Binomial logistic regression Model 2, reduced from the full, main effects model (Table S2, Model S4), used to explain the effects of sex on the presence of *Hematodinium* spp. in 2011.

### Injury

Comparative analysis between the two sites (fished; RZ vs. un-fished; NTZ) revealed that there was no significant change in injury prevalence between sites. However, when combining data from both sites, analysis highlighted that the percentage of injured crabs (examples of which can be seen in Fig. 2E-G) increased significantly by *ca*. 12% between 2010 and 2011 (P < 0.001, Table 2).

In 2010, landing size, site and sex were not significant in predicting the presence of injury (Table S2, Model 5), even after model reduction, and therefore no reduced model was produced. In 2011, although none of the predictor variables significantly predicted the presence of injury in the full model (Table S2, Model S6) after model reduction, landing size was significant in predicting injury presence (Table 5). Crabs below the MLS in 2011 had an 11% higher probability of being injured than those above the MLS (Fig. 7).

**Table 5.** Binomial logistic regression Model 3, reduced from the full, main effects model (Table S2, Model S6), used to explain the effects of landing size on the presence of injury in 2011.

| Model | Predictor variable | Estimate<br>(slope) | <sup>±</sup><br>Standard<br>Error | P value |
| --- | --- | --- | --- | --- |
| *Injury ~ Landing Size* |  |  |  |  |
| *df = 182* | *Landing size*<br>*(>MLS)* | -2.0458 | 1.0366 | 0.0484 * |
| *AIC: 150.38* |  |  |  |  |

\* Asterisk denotes significance ( $\alpha \leq 0.05$ ).

## DISCUSSION

Overall, more crabs were caught in the RZ compared with the NTZ in both years, rejecting our first hypothesis. Our second hypothesis was also rejected (crabs from the un-fished NTZ would have an increased probability of injury and therefore disease than individuals from the fished RZ). Shell disease increased from 2010 to 2011 in line with injury. However, only if a crab was male or missing a limb, did the probability of it having shell disease increase in 2011. Crabs below the MLS in 2011 had a higher probability of being injured than those above the MLS, rejecting the third hypothesis that larger animals would have higher disease/injury.

### Hypothesis 1

The population ecology observed in this study contrasted to that of a previous survey of *C. pagurus* around Lundy Island MPA. In a 4-year survey from 2003-2007, Hoskin et al. (2011) described higher abundances, and larger *C. pagurus* within the NTZ than the RZ. This contrasts to the current results from 2010 and 2011 in which we found higher abundances of *C. pagurus* in the RZ. There are various possible explanations for this finding. *C. pagurus* migrate up to 345 m day^−1^ in order to avoid predators, competition from conspecifics, and to mate or find brooding sites (Ungfors et al. 2007, Hunter et al. 2013), it is therefore plausible that the low crab abundance described in 2011 is due to natural movement of populations out of the NTZ into adjacent areas. However, in addition, density dependent habitat selection whereby an increased abundance of one species may change the dynamics of another has been described in detail by many authors (e.g. Breen & Mann 1976, Morris 2003, Acheson & Gardener 2014). European lobsters, *H. gammarus*, were also caught in the pots sampled during this study (further discussed in Davies et al. 2015) and it is noteworthy that as the abundance of lobsters per string increased, the abundance of crabs decreased significantly (2010: R^2^ = 0.537, F_1, 14_ = 16.22, P = 0.0012, 2011: R^2^ = 0.437, F_1,12_ = 9.320, P = 0.01; Fig. 4). Although studies have shown that if a larger predator is present in a pot, other animals will not enter as willingly (Lovewell et al. 1988, Miller & Addison 1995), there is also evidence to that low *C. pagurus* abundances may be driven by increased abundances of the European lobster, *Homarus gammarus,* found in the NTZ (Wootton et al 2012, Davies et al. 2015). This phenomenon was also observed by Howarth et al. (2016) who noted that in a fully protected MPA in Scotland, the greater densities of large adult lobsters appeared to be predating and/or competitively displacing juvenile lobsters, *C. pagurus*, and velvet swimming crabs (*Necora puber*) from the area. This highlights community changes as a result of MPA implementation which may not always be positive, and that recovery is not straightforward, since the recovery of some species can have knock-on effects on others. As one species benefits from implementation (in this case, the apex predator, *H. gammarus*), others (i.e. *C. pagaurus*) can be detrimentally affected, altering overall ecosystem function.

**Figure 4.**
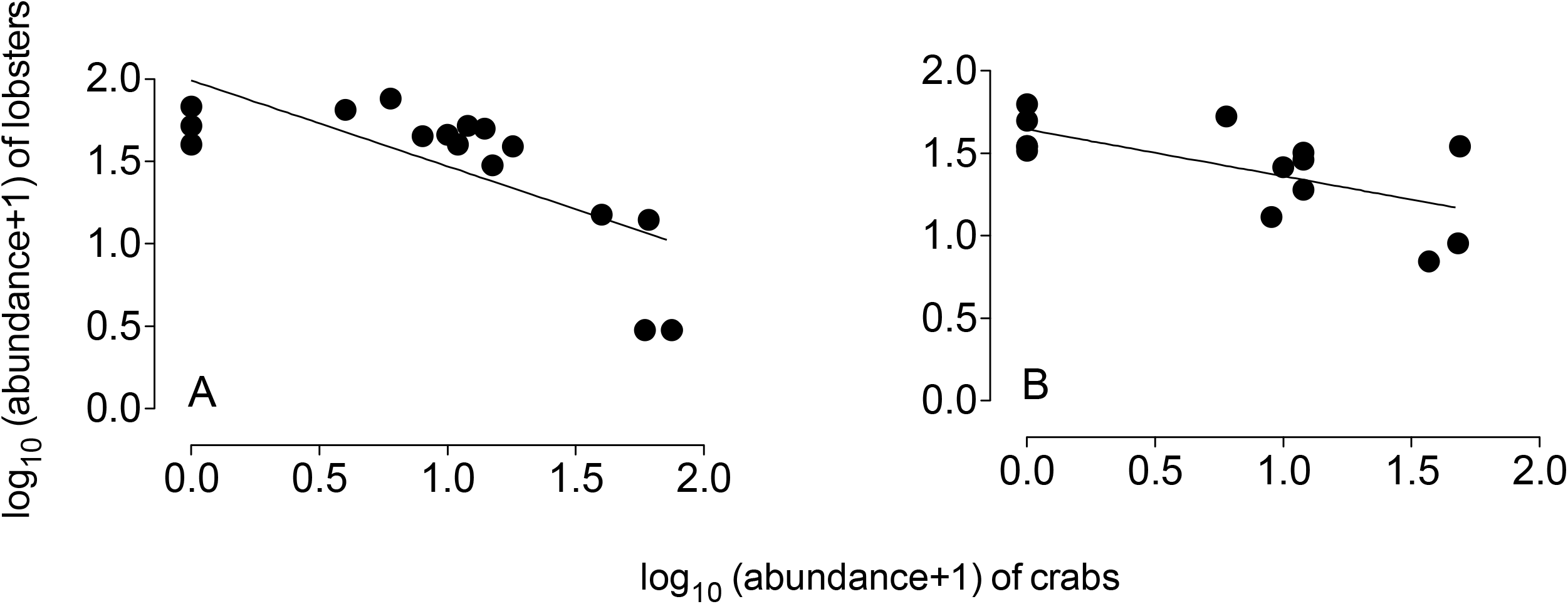
Linear regression showing the abundance of lobsters plotted against the abundance of crabs per string in (A) 2010 and (B) 2011. Each point represents one string of 35 parlour pots.

**Figure 5.**
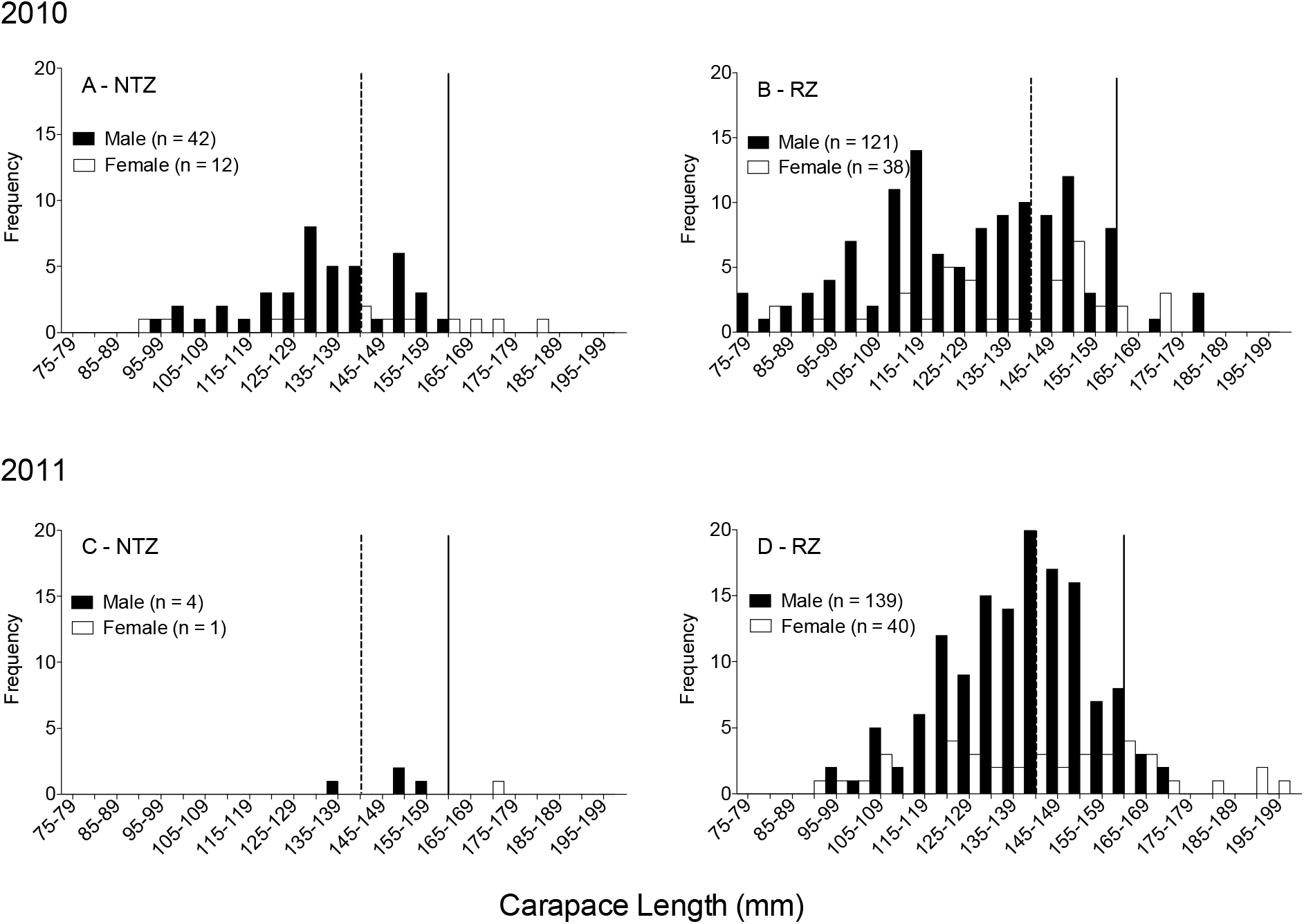
Size-frequency distributions of male and female crabs surveyed from 2010 (A, B) and 2011 (C, D). Broken lines indicate minimum landing size (MLS) for females (carapace width ≥140 mm) and solid lines indicate MLS for males (carapace width ≥ 160 mm).

**Figure 6.**
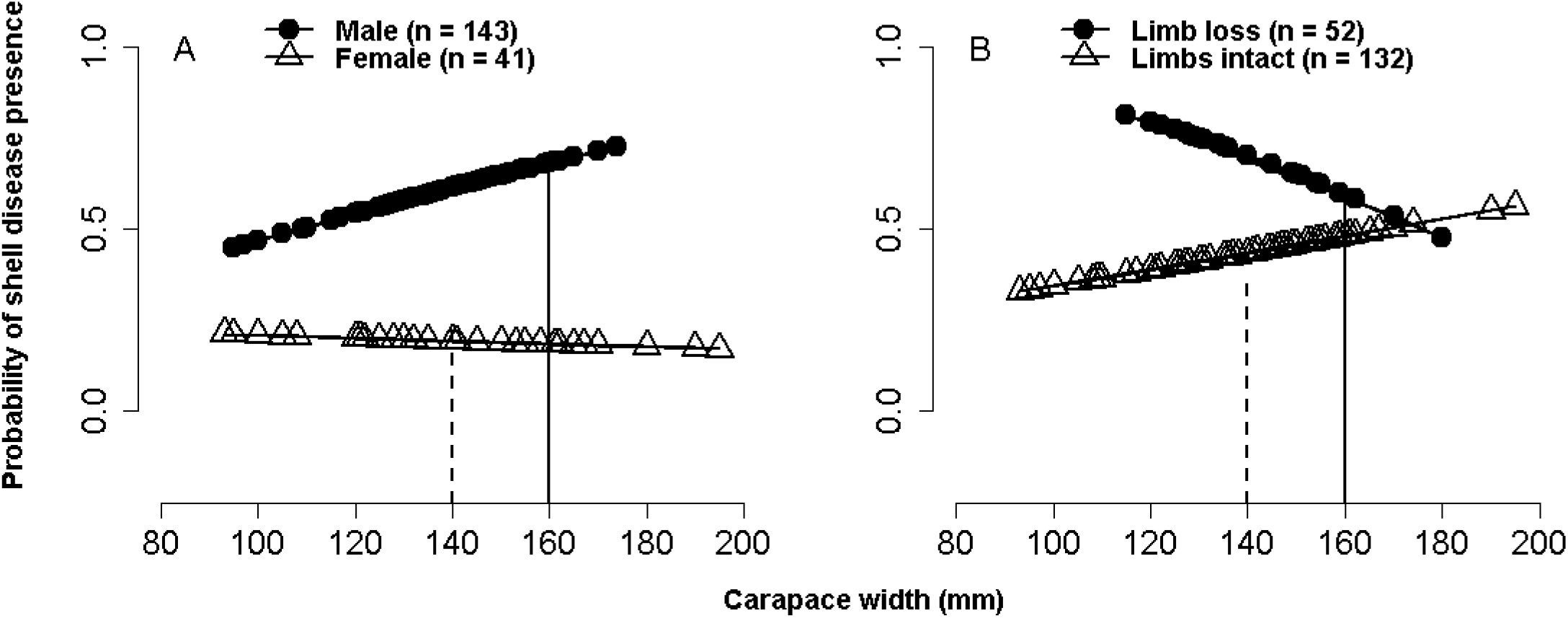
Fitted probability plots of shell disease presence in 2011 against carapace width, separated by significant predictor variables (A) sex and (B) limb loss (Table 3). The broken line in each plot indicates minimum landing size (MLS) for females (carapace width ≥140 mm) and the solid line indicates MLS for males (carapace width ≥ 160 mm).

**Figure 7.**
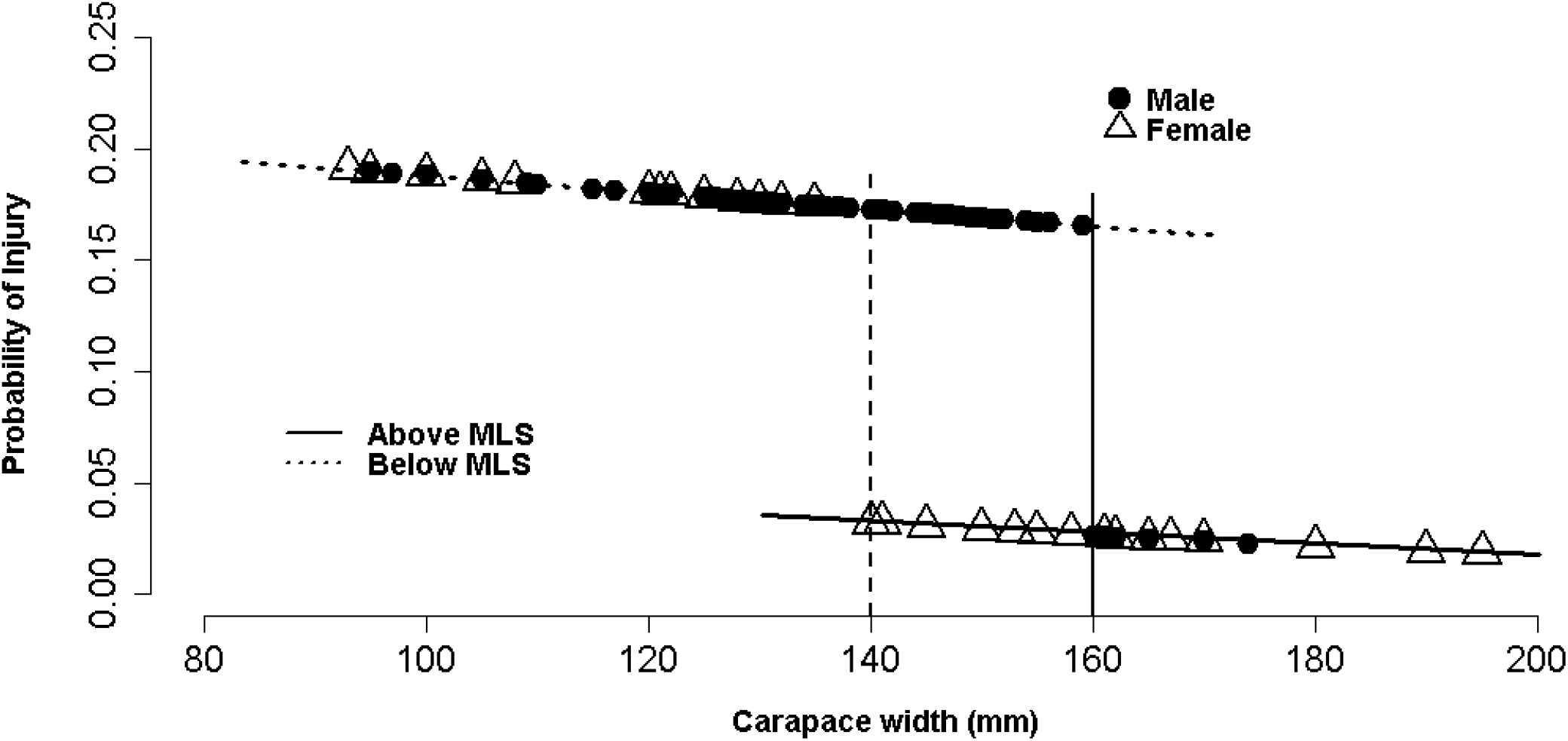
Fitted probability plot of injury presence against carapace width, separated by the significant predictor variable of above *vs.* below minimum landing size (MLS) (Table 5). The broken line indicates minimum landing size for females (carapace width ≥140 mm) and the solid line indicates MLS for males (carapace width ≥ 160 mm).

### Hypothesis 2

Crabs were no more likely to be found with injury, limb loss or disease in the un-fished NTZ compared to the fished RZ as has been found in previous studies. This is likely due to higher population density in the RZ and interactions with other species. The higher levels of shell disease found in males compared to females (in both sites combined) is likely caused by increased male-male interaction and resultant injuries during the mating season (the time of our study). Male crabs engage in agonistic behaviour with conspecifics due to their territorial and competitive nature, increasing the risk of injury (Schöne 1968, Vogan et al. 1999) limb loss and therefore disease (Vogan et al. 2008). Lost limbs create a large aperture for haemolymph loss, tissue exposure and pathogen entry, in addition to an increased risk of shell disease lesion initiation stemming from the fracture site. This may therefore account for shell disease presence being higher in those crabs that had lost one or more limbs.

### Hypothesis 3

The unfished NTZ in Lundy was not found to harbour larger crabs than in the fished, RZ as previously noted by Hoskin et al. (2011). Similarly, larger crabs (>MLS) were not found to exhibit more injury and disease as previously found in some crustacean species (Bateman et al. 2011, Wootton et al. 2012, Davies et al. 2015). In contrast, the probability of injury increased significantly in both male and female crabs **below** the MLS. *C. pagurus* above the MLS are sexually mature and potential injury above this size in theory is likely to increase due to conflict from protection and display during mating (Edwards 1966). However, one explanation for the increased injury in individuals below the MLS found in the present study, could be due to competitive interactions with the significantly more abundant, larger and more aggressive predator, the lobster, *H. gammarus*. Another explanation could be injury from species on which *C. pagurus* feeds. *C. pagurus* is an active predator that consumes a variety of crustaceans including the green shore crab *Carcinus maenas*, which is an aggressive and territorial species (Kaiser et al. 1990). *C. maenas* has been shown to contribute to injury in its predators especially if the predators are smaller and inexperienced hunters (Rossong et al. 2006), and this could include small *C. pagurus* (below the MLS).

Overall, there were very few crabs caught infected with *Hematodinium* spp. over both years sampled. Similarly, low numbers have been described in other studies of wild *C. pagurus* populations (Chualáin et al. 2009) and may be explained by the observation that *Hematodinium* spp. in *C. pagurus* is mainly found in smaller or juvenile crabs which live closer to the shoreline than the areas sampled in this study (Stentiford 2008, Chualáin et al. 2009). Therefore, most crabs surveyed in the current study were adults (even though escape gaps on sampling pots were closed). Due to the low numbers of *Hematodinium* spp. infected individuals found in this study, it would be beneficial in future studies to survey closer to shorelines and modify traps to restrict larger crab and lobster entry. This would allow for a better census of juveniles, who are more likely to display the disease.

The current study only provides a ‘snap shot’ of population abundance and disease ecology. It does, however, reiterate the importance of disease monitoring within both fished and protected populations, and especially those that have shown significant increases in local abundance. Long-term monitoring studies of MPAs (those observed over a number of years) have revealed strong patterns, including spillover into adjacent fisheries and significant increases in abundance (Abesamis & Russ 2005, Stobart et al. 2009, Aburto-Oropeza et al. 2011) that may not be possible to detect during small-scale studies such as this. It would therefore be beneficial to monitor MPAs such as Lundy for longer periods, accounting for natural variability driven by external factors such as adjacent fisheries, natural competition and climate anomalies. At present, disease monitoring does not appear to be a priority in the implementation and management of protected areas. By monitoring disease, managers can be better prepared to deal with any unwanted consequences of fisheries closures such as potential increases in disease and consequent population decline. Better monitoring will allow pre-emptive management measures to be taken. An example of this could be the re-opening of closed areas for limited fishing of certain species, before levels of abundance become high enough to contribute to disease. This would allow real-scale tests of ‘fishing out’ disease, or help strike a balance between allowing enough fishing to keep populations healthy without interfering with explicit conservation aims (McCallum et al. 2005, Wood et al. 2010).

To effectively and sustainably manage, exploit and conserve marine populations, it is imperative to monitor both the prevalence and geographical range of important marine pathogens, especially those affecting keystone and species of commercial interest (McCallum & Dobson 1995, Lamb et al. 2016). This is particularly pertinent for protected areas, where detrimental secondary community effects have been shown to occur. Robust monitoring programmes in such areas, covering a range of species and variables, would assist in achieving conservation aims and allow management strategies to be adjusted according to local ecological change.

## Acknowledgements

This research was funded by SEAFISH and the ERDF INTERREG IVA, Ireland-Wales programme grant -SUSFISH (Project No. 042) to AFR. CED was part-funded by a tuition fee bursary from Swansea University’s Colleges of Science and Medicine and would also like to thank the Society of Biology and Climate Change Consortium for Wales (C3W) for travel grants enabling the collaboration with the AVC Lobster Science Centre. The authors would like to thank Geoff Huelin and the crew of FV ‘Our Jenny’, plus Devon and Severn IFCA and Natural England for permission to sample crabs in the No-take Zone of Lundy Island. We also thank Dr. Gethin Thomas, Dr. Richard Unsworth, Dr. Ed Pope, Dr. Kristina Hamilton, Keith Naylor and Ian Tew of Swansea University, and Lundy Island staff for their support during sampling. We also thank Mrs Carolyn Greig of Swansea University and Mr Adam Acorn of the AVC Lobster Science Centre, UPEI for laboratory assistance and Dr Hilmar Hinz of IMEDEA, Mallorca for statistical advice.

