## Supplementary Materials for "Factors contributing to the disease ecology of brown crab (*Cancer pagurus*) in a temperate marine protected area"

**S1. Modified BCD version of Ivanova et al. (2006) 96 well Plate Extraction Protocol**

***Pre-DNA Extraction:***

Note: The night before extraction: check wells to see if the EtOH has evaporated and if so then top them up to original 1ml volume. Place sample plate on its side in a shaking incubator overnight (shake, no heat). ***Add a known positive blood sample to B10 of the plate.**

***Day 1:***

1. Vortex plate and transfer 150 µl blood-EtOH into 0.2ml strip tubes for lysis. (**Cut ends off of tips to assist in pipetting blood**)
2. Cover tubes and centrifuge for 3 min at 2000g to pellet blood. Removing only one strip of tubes at a time, pour off ethanol onto thick paper towel. Let the residue evaporate for 30 min to 1 hour in the 56°C incubator
3. Recap original Sample plate with new caps to prevent cross contamination.
4. Combine 5ml of Insect Lysis Buffer per plate and 0.5ml of Proteinase K (20mg/ml) in a sterile container. Add 50 µl of lysis mix to each well of blood. Cap and vigorously vortex to mix in lysis mix.
5. Incubate at 56°C overnight (24 hour incubation gives better yield)

***Day 2:***

1. Centrifuge at 1500 g for 2 min to remove condensation from strip caps.
2. Add 100 µl of Binding Mix to each sample and shake vigorously for 10-15 sec. Centrifuge at 1000g for 2 min.
3. Transfer lysate (about 150 µl) to the wells of the 96 well filter plate (No. 8032; Pall Life Sciences) on top of the square-well block and seal plate.
4. Centrifuge at 6000 g for 7 min to bind DNA.
5. First Wash: Add 180 µl of Protein Wash Buffer to each well and seal with a new cover. Centrifuge at 6000 g for 5 min.
6. Second Wash **(2X)**: Add 750 µl of Wash Buffer to each well and seal with new cover. Centrifuge at 6000 g for 7 min. **Repeat this step twice.**
7. Open sealing cover, close it and centrifuge at 6000 g for 5 min to completely remove the Wash Buffer.
8. Remove cover and place plate on the lid of a tip box. Incubate at 56°C for 30 min to evaporate residual EtOH. **Aliquot nuclease free water into 15 ml sterile tube. Aliquot 10xTE buffer into 1.5 ml sterile tube. Place both aliquots in incubator with plates.**
9. Position PALL 2 collar on the collection microplate and place the filter plate on top. Dispense 50 µl of preheated (56°C) ddH_2_O onto the membrane of each well and incubate at RT for 2 min. Seal plate.
10. Place assembled plates on a clean square-well block to prevent cracking and centrifuge at 6000 g for 5 min to collect DNA elute. Remove and discard PALL plate.
11. Add 5.5 µl of 10X TE buffer to each sample. Label plate with Sample plate ID, date and Initials both on plate and foil. Proceed with PCR reaction or cover plate and store at -20°C.

***Stock Solutions:***

Solution	Components	Final Volume

**1M Tris-HCl, pH 8.0:**	Tris base 60.57 g	500 ml

HCl

**1M Tris-HCl, pH 7.4:**	Tris base 60.57 g	500 ml

HCl

**0.1M Tris-HCl, pH6.4:**	Tris base 6.06 g	500 ml

(adjust pH with HCL to 6.4-6.5)

**1M NaCl:**	NaCl 29.22 g	500 ml

**0.5M EDTA pH 8.0:**	EDTA 93.05 g	500 ml

NaOH ~10.0 g

(NOTE: Vigorously mix on a stirrer with heat. The disodium salt will not go into sol’n until the pH is approx. 8.0 by the addition of NaOH).

**Proteinase K:**	100mg	5 ml

Reconstitute to a final volume of 5 ml in Molecular grade Nuclease Free water

Aliquot by 0.5 ml. Store at -20°C.

- Wash all labware with ELIMINase and rinse with ddH_2_0. Filter buffers through 0.2 µm filter into a clean bottle. Make smaller working volume aliquots (eg. 100 ml). Store stock solutions and working aliquots at 4°C, unless otherwise stated.

***Working Solutions for DNA Extraction:***

Buffer	Vol. stock	Initial stock conc.	Final Volume

**Insect Lysis Buffer:**

700mM GuSCN	16.5g	200 ml

30mM EDTA	12 ml	**0.5M EDTA, pH 8.0**

30mM Tris-HCl	6 ml	**1M Tris-HCl, pH 8.0**

0.5% Triton X-100	1 ml

5% TWEEN 20	10 ml

NOTE: Vigorously mix on a stirrer with heat.

**Binding Buffer:**

6M GuSCN	354.6 g	500 ml

20mM EDTA	20 ml	**0.5M EDTA, pH 8.0**

10mM Tris-HCl	50 ml	**0.1M Tis-HCl, pH 6.4**

4% Triton X	20 ml

NOTE: Vigorously mix on a stirrer with heat. If any recrystalization occurs, then pre-warm at 56°C to dissolve before use.

**Wash Buffer:**

60% etOH	300 ml	**100% etOH**	500 ml

50 mM NaCl	25 ml	**1M NaCl**

10mM Tris-HCl	5 ml	**1M Tris-HCl, pH 7.4**

0.5mM EDTA	0.5 ml	**0.5M EDTA, pH 8.0**

Sterile water	169.5 ml

NOTE: mix well, store at -20°C.

**Binding Mix:**

Binding buffer	50 ml	100 ml

100% etOH	48 ml

Sterile water	2 ml

**Protein Wash Buffer:**

Binding buffer	26 ml	100 ml

100% etOH	67.2 ml

Sterile water	6.8 ml

NOTE: Stable at room temperature for about 1 week, discard if crystallization occurs.

*Make sure that there are two empty wells left for the controls; discard any liquids that may have eluted into the wells.

**S2. Decapod PCR conditions**

MangoMix (Bioline Ltd, UK) and decapod-specific primers (143F 5'-TGCCTTATCAGCTNTCGATTGTAG-3' and 145R 5'-TTCAGNTTTGCAACCATACTTCCC-3' yielding an 848 bp amplicon; (N represents G, A, T, or C) were used to verify the quality of the extracted DNA and the integrity of the PCR reaction (Lo 2014). Cycling conditions were as follows: 4 min at 94°C followed by 40 cycles of 1 min at 93°C, 1 min at 55°C and 2 min at 72°C, followed by 5 min at 72°C.

**S3. Hematodinium spp. specific PCR conditions**

DNA was amplified in 10 μ1 total reaction volume by adding 2 μl of 5x Colored Reaction Buffer, 375 µM of each dNTP, 3 mM MgCl_2_, 2 µM of each primer, 0.05 μl of MangoTaq polymerase (1U/µl, Bioline Ltd, UK), 2 μl of diluted DNA (1:10), 10% BSA and sterile water to the final reaction volume. *Hematodinium* spp. specific primers optimized by Hamilton *et al.* (2009) were used: DinoF 5'-GTGGTGCATGGCCGTTCTTAGTT-3 and ITS1R1 5'-GAAGGGAAGGGGAGAAGAAGC-3. Cycling conditions were as follows: 3 min at 95°C followed by 34 cycles of 45 sec at 94°C, 45 sec at 57°C and 1 min at 72°C, followed by 10 min at 72°C.

**S4. European lobsters**

European lobsters, *H. gammarus*, were also caught in the pots sampled (see Davies *et al.* 2015). A linear regression was used to examine the relationship between the CPUE of *C. pagurus* and European lobster, *H. gammarus*, the two commercially viable species found in the pots. It is noteworthy that as the abundance of lobsters per string increased, the abundance of crabs decreased significantly (2010: R^2^ = 0.537, F_1, 14_ = 16.22, p = 0.001, 2011: R^2^ = 0.437, F_1,12_ = 9.320, p = 0.01; Fig. S1).

**Table S1.** **Details of data collection**

| **Year** | 2010 |  | 2011 |
| --- | --- | --- | --- |
| **Month** | May | July | August |
| **Total crabs caught** | 165 | 48 | 184 |
| **Total crabs removed (as day recaptures)** | 2 | 1 | 2 |

**Table S2.** Full models used in order to predict response variables of shell disease, *Hematodinium* spp. and injury before reduction. P values were calculated using an ANOVA of the null versus the residual model, pseudo R2 are calculated using deviance measures ((null-res)/null). Asterisk denotes significance (α ≤ 0.05).

| **Model** | **Parameter** | **Estimate**  **(slope)** | **p** |
| --- | --- | --- | --- |
| Model S1: Shell disease full model 2010 (main effects)  Shell Disease ~ Injury * Landing size + Limb loss + *Hematodinium* + Sex + Site  **AIC:** 284.17 | Injury  Landing size  Limb loss  *Hematodinium*  Sex  Site | -0.65  -0.78  -0.67  0.04  0.78  -0.12 | 0.579  0.866  0.862  0.946  0.068  0.725 |
| Model S2: Shell disease full model 2011 (main effects)  Shell Disease ~ Sex * *Hematodinium* + Limb loss + Injury + Landing size + Site  **AIC:** 244.12  Model S3: *Hematodinium* full model 2010 (main effects)  *Hematodinium* ~ Site * Limb loss + Injury + Landing size + Sex  **AIC:** 94.14  Model S4: *Hematodinium* full model 2011(main effects)  *Hematodinium* ~ Limb loss + Injury + Landing size + Sex + Site  **AIC:** 67.63  Model S5: Injury full model 2010 (main effects)  Injury ~ Landing size + Sex + Site  **AIC:** 56.94  Model S6: Injury full model 2011 (main effects)  Injury ~ Landing size + Sex + Site  **AIC:** 154.20 | Sex  *Hematodinium*  Limb loss  Injury  Landing size  Site  Site  Limb loss  Injury  Landing size  Sex  Limb loss  Injury  Landing size  Sex  Site  Landing size  Sex  Site  Landing size  Sex  Site | 1.99  1.85  1.04  -0.05  0.11  1.56  -0.13  -0.41  -0.30  -15.24  1.00  1.19  -16.81  -1.78  -2.32  -16.28  1.30  1.10  0.45  -1.66  0.83  0.34 | <0.001*  0.098  0.008*  0.920  0.823  0.220  0.879  0.711  0.804  0.992  0.401  0.189  0.992  0.133  0.012*  0.977  0.192  0.375  0.686  0.127  0.302  0.771 |
